# Data science approaches provide a roadmap to understanding the role of abscisic acid in defence

**DOI:** 10.1101/2022.05.30.493976

**Authors:** Katie Stevens, Iain. G. Johnston, Estrella Luna

**Affiliations:** School of Biosciences, University of Birmingham, Birmingham, B15 2TT, UK; Department of Mathematics, University of Bergen, Norway; Computational Biology Unit, University of Bergen, Norway

**Keywords:** ABA, plant hormone, resistance, decision tree, machine learning

## Abstract

Abscisic acid is a plant hormone well known to regulate abiotic stress responses. ABA is also recognised for its role in biotic defence, but there is currently a lack of consensus on whether it plays a positive or negative role. Here, we used supervised machine learning to analyse experimental observations on ABA to identify the most influential factors determining disease phenotypes. ABA concentration, plant age and pathogen lifestyle were identified in our computational predictions. We explored these predictions with new experiments in tomato, demonstrating that phenotypes after ABA treatment were highly dependent on plant age and pathogen lifestyle. Integration of these new results into the statistical framework refined the quantitative model of ABA influence, suggesting a framework for proposing and exploiting further research to make more progress on this complex question. Our approach provides a unifying road map to guide future studies involving the role of ABA in defence.

## Introduction

Abiotic and biotic defence responses are important evolutionary mechanisms that allow plants to adapt to their dynamic environments (Peck and Mittler, 2020). A variety of sophisticated signalling networks modulate plant responses to challenges, helping to enhance survival. Hormone signalling and crosstalk play major roles in these responses, which can also be influenced by plant age, organ type and environment (Berens et al., 2017). However, as plants encounter many biotic challenges, including pathogens with varying lifestyles and methods of attack, there is no one-size-fits-all approach to defence. Therefore, hormones have unique roles in multiple, often antagonistic, complex defence pathways (Robert-Seilaniantz et al., 2011).

Abscisic acid is an essential hormone involved in both biotic and abiotic stress signalling (Lee and Luan, 2012). ABA is found universally across vascular plants and is involved in a wide range of physiological processes. For instance, through interaction with developmental hormones, ABA modulates plant growth, seed germination, stomata closure and embryo maturation (Vishwakarma et al., 2017). Moreover, ABA is essential for adaptation to drought and salt stress (Sah et al., 2016) and has a positive role in ensuring plant survival during abiotic stress. For example, water supply limitation is associated with an increase in ABA across the entire plant that leads to reduced stomatal conductance (Vishwakarma et al 2017). However, the function of ABA in defence against biotic challenges, including pathogens, is varied, which has led to mounting controversy in the field of plant biology (Asselbergh et al., 2008a, Cao et al., 2011, Ton et al., 2009).

ABA is widely considered to be a positive influence on early pathogen invasion due to its essential role in stomatal closure (Melotto et al., 2006). However, controversy arises in the later infection process. On one hand, ABA is known to be involved in multiple defence pathways and often interacts antagonistically with other hormones such as salicylic acid (SA), jasmonic acid (JA) and ethylene (ET) through ‘hormone crosstalk’ (Anderson et al., 2004, De Torres Zabala et al., 2009, Oide et al., 2013). SA is involved in resistance against biotrophs and its production is enhanced following colonisation with the bacterial biotroph *Pseudomonas syringae* (Delaney et al., 1995). However, SA has a negative role in resistance to necrotrophs such as *Botrytis cinerea* (El Oirdi et al., 2011). In contrast, JA and ET display the reverse effect of SA and enhance resistance against necrotrophs and herbivory (McDowell and Dangl, 2000). These hormones therefore behave antagonistically and ABA is typically shown to have a negative influence on resistance as it can disrupt SA-, JA-and ET-mediated defence signalling (Anderson et al., 2004, Audenaert et al., 2002). On the other hand, ABA has been linked to the expression of defence responses controlled by these plant hormones, including deposition of callose and reactive oxygen species production, which mediate resistance against biotrophic and necrotrophic pathogens (Asselbergh et al., 2007). In addition, crucially, a positive role of ABA in defence has been linked to the requirement of an intact ABA signalling pathway for direct and priming of callose deposition (Flors et al., 2005, Flors et al., 2008, Luna et al., 2011, Schwarzenbacher et al., 2020, Ton and Mauch-Mani, 2004, Vicedo et al., 2009). Therefore, the role of ABA on the expression of defence mechanisms appears highly varied, with positive and negative effects via different mechanisms during different circumstances.

The effect of ABA does not appear to be associated with plant species or pathogen lifestyle (Asselbergh et al., 2008b), and contradictions have been shown even within a single model plant pathosystem. For example, whereas ABA-treated Arabidopsis plants display enhanced susceptibility against *P. syringae* (Melotto et al., 2006, Mohr and Cahill, 2007), Melotto et al showed that the ABA-deficient *aba3-1* Arabidopsis mutant displays enhanced susceptibility against *Pst*DC3000 (Melotto et al., 2006). In economically important crops such as tomatoes, ABA deficient *sitiens* tomato mutants have enhanced resistance against *B. cinerea, Odium neolycopersici* and *Erwinia chrysanthemi* (Achuo et al., 2006, Asselbergh et al., 2007, Asselbergh et al., 2008b, Audenaert et al., 2002). In contrast, long-lasting induced resistance in tomato fruit is marked by an accumulation of ABA (Wilkinson et al., 2018). ABA is therefore a key component of defence at all phases of infection, yet its influence remains misunderstood.

To make progress understanding such complex questions, machine learning and supervised learning techniques are increasingly being used for studying plant pathogen interactions, specially motivated by the increased production of large -omics datasets and imaging capabilities (Sperschneider, 2020, Sun et al., 2020, van Dijk et al., 2021). For instance, ML has facilitated analysis of gene networks in immune response (Dong et al., 2015), pathogen effector protein localisation (Sperschneider et al., 2018) and plant disease diagnostics using image-based learning (Mishra et al., 2019). Here, we have used ML strategies to investigate the contrasting effects of ABA in resistance against pathogens. Through the collection of hundreds of data points from a total of 30 scientific peer-reviewed publications, we used tools from ML to attempt to unpick the complex interplay of effects shaping heterogeneous ABA influences, combined with new experiments to further explore these data-driven findings. Overall, our study highlights key factors that affect the effect of ABA in resistance and provides a tool for future studies towards facilitating the understanding of the role of ABA in plant responses against biotic stresses.

## Methods

### Data collection and preparation

Quantitative data regarding disease resistance was collected from literature involving ABA (Table S1). A total of 30 peer reviewed papers were selected after a broad literature search as they included quantitative data from experiments directly involving pathogen infection and ABA including ABA signalling and biosynthesis mutants, exogenous application of ABA, and/or ABA inhibitors. From those papers, 13 shared variables were recorded from the three general classes of plant, ABA treatment, and pathogen. From the plant, we recorded the species, variety, genotype, mutant, mutant type, plant age and tissue. From ABA, we recorded the treatment, application method, and concentration applied. From the pathogen, we recorded the type, lifestyle, and species. All variable information was taken directly from papers used, however values of plant age for orange, grape and pepper were imputed based on known cultivar information. A large number of disease scoring methods were used across publications such as lesion diameter (mm), % spreading lesions, and categorical scoring. A disease severity index (DSI) was therefore used in an attempt to unify the heterogeneity of disease scoring in our dataset. All observations involving ABA were compared to the level of resistance in control treatments for each individual publication. DSI was represented as a ‘percentage resistance change’ of all data points, forming a continuous measure of percentage resistance change as a result of ABA, as described below (Table S1, Figure 1).

**Figure 1:**
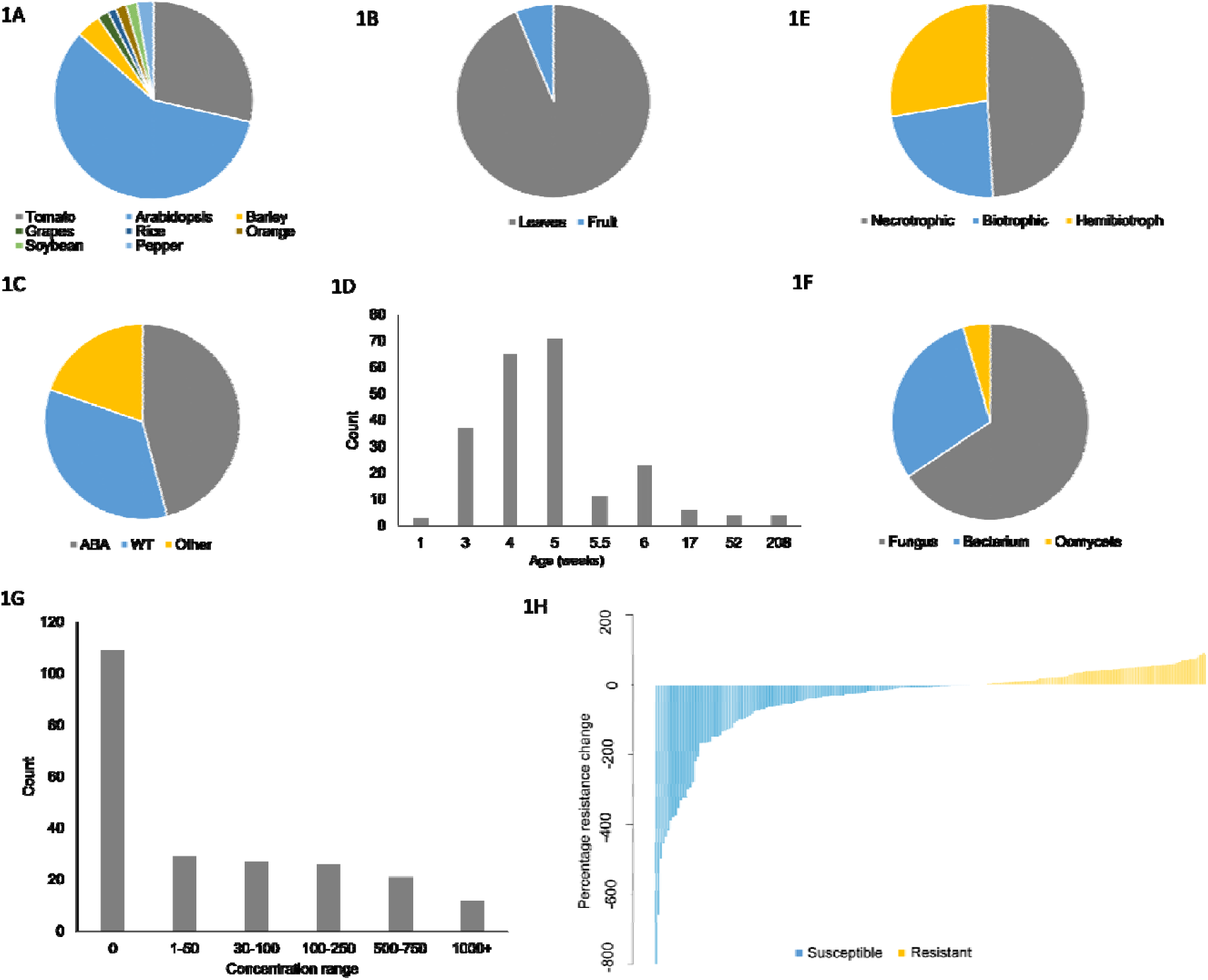
Characteristics of our amalgamated dataset. Proportion of plant species (A), plant tissue (B), mutant type (C), count distribution of plant ages (weeks) (D), pathogen lifestyle (E), pathogen type (F), and count distribution of ABA concentration range (G). (H) Distribution of percentage resistance value. Data described in Table S1.

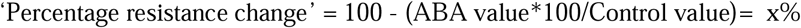

### Computational meta-analysis

Two supervised learning approaches, described below, were used to identify the most important factors to predict disease resistance (Breiman et al., 1984). All analysis was conducted in R (Version 1.4.1717). For each method of classification, data was randomly split into training (75%) and test (25%) data. Models were generated using the training data and model accuracy was assessed using test data. Final models were generated using all data available. Two models were created: a decision tree (DT) predicting a binary response of resistance ‘susceptible’ or ‘resistant’, depending on if the percentage resistance change was a positive or negative value, and a random forest model. The same variables and parameters were used for both DT and RF. To decide which variables to include in our models and guard against overfitting, the Akaike Information Criterion (AIC) score was used, providing a trade-off between the (maximum likelihood) fit of the model on the training data set and the complexity of the model, reflected by the number of parameters. The model with the lowest AIC score was selected.

### Classification and regression tree (CART)

Decision trees (DT) were constructed using rpart (v 4.1-15) (Therneau et al., 2015) with the data described in Table S1 and Table S2. Rpart uses the classification and regression tree (CART) algorithm to construct trees based on recursive binary partitioning, which splits the data based on the best predictive variable. The chosen variables and specific splits are determined by the Gini impurity index which measures the disorder in a set of data points. Gini impurity is calculated as the probability of incorrectly labelling a data point assuming random labelling according to the distribution of all classes in the set (Breiman et al., 1984). Splits in trees therefore aim to maximise the decrease in impurity; this process is repeated to build the tree. Seven variables were selected for DT analysis based on the optimal AIC score. Trees produced via the CART algorithm are typically large and overfitted, to reduce complexity trees are pruned. Pruning is determined by the rpart ‘complexity parameter’ value of the smallest tree with the smallest cross validation error (Therneau et al., 2015). The CART algorithm additionally provides a measure of variable importance. Variable importance is determined by the accumulated contribution of each variable across all nodes and trees where it is used (Breiman, 2001, Breiman et al., 1984).

### Random forest

Random forest analysis was performed using the package randomforest (v 4.6-14) (Liaw and Wiener, 2002) using the data described in Table S3 and Table S4. RF uses the CART algorithm, as described above, to construct a set of different individual trees, which are assembled into a ‘forest’. Here, each tree provides a classification ‘vote’, and the average vote of all trees in the forest is selected. Through resampling with replacement, random forests guard against overfitting, and RFs can improve accuracy and predictive abilities in classification compared to singular CART trees (Breiman, 2001). In our approach, the number of trees and cross validation was optimised to provide the lowest out-of bag (OOB) estimate of error rate. The final RF model used in this study contained 400 trees and 10 folds cross-validation.

### Plant materials and growth conditions

Tomato seeds (cultivar micro-tom) were maintained in damp, humid, dark conditions to stimulate germination. Germinated seeds were transferred to individual 80mL pots containing Scott’s Levington M3 soil. For analysis of older plants, three-week-old plants were transplanted into 3 L pots until experiments were performed. Plants were grown in greenhouses under 16/8-hour day/night cycles, 150 µmol m^-2^ s^-1^ light intensity and 25°C /20°C temperatures. Experiments were performed between July and November 2020.

### Abscisic acid treatment

Leaf material was treated with freshly prepared Abscisic Acid (Sigma Catalogue number A1049). ABA was dissolved in ethanol and then diluted in water to the different concentrations used in this study: 20_μ_M, 50_μ_M, 100_μ_M, 500_μ_M. The control treatment was water. All treatment solutions were adjusted to the amount of ethanol present, determined by the highest concentration of ABA. Leaves from 4-weeks and 8-weeks old plants were excised and placed in a plastic tray, lying flat, with stems wrapped in wet paper as previously described (Luna et al., 2015, Worrall et al., 2012). Two leaves per plant and a total of six plants (six biological replicates) were used to test the effect of ABA in leaves. ABA treatment was performed by spraying the treatment solutions onto leaves. Leaves were then allowed to dry before moving trays to 100% humidity in the dark for 24h before pathogen infection.

#### *Botrytis cinerea* cultivation and infection

Infections of leaves with *Botrytis cinerea* (isolate BcI16) were performed as previously described (Luna et al., 2015) infected with droplets of 5_μ_L inoculum containing 5×10^5^ spores/ml. Leaves were incubated in the dark at 100% humidity and 20°C. Disease severity was recorded at 3-days post infection (dpi) by measuring the diameter of the necrotic lesions using a Vernier calliper.

#### Phytophthora infestans cultivation and infection

*Phytophthora infestans* 88069td10 was grown on rye medium supplemented with geneticin for 2-3 weeks at 20°C as previously described (Whisson et al., 2007). Plates were flooded with 10ml water and spores scraped from the plate surface and incubated at 4°C for 1-3 hours. Zoospores released from sporangia were decanted. A desired concentration of 5×10^4^ spores/ml was obtained. The underside of each leaf was infected with six droplets of 10_μ_l inoculum. Plants were incubated at 100% humidity, 16hour/8hour light/dark cycles and 18°C /15°C temperature cycle. Disease severity was recorded at 7-days post infection. Scoring was done by classifying lesions into different categories of colonisation. Class I: healthy, Class II; lesions on leaf underside only, Class III; lesions in under and upper leaf surface, Class IV: prominent lesions on both leaf sides, Class V: spreading lesions causing tissue damage, Class VI: total leaf collapse. Category distributions were used to calculate disease severity indexes using the formula (Chiang et al., 2017):

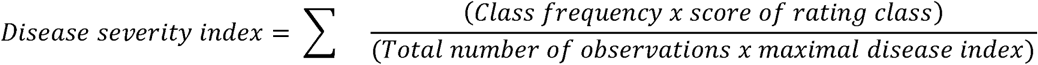

### Statistical analysis

Data from the experiments with *B. cinerea* and *P. infestans* were analysed using R (v 1.2.1335). In the analysis of lesion diameter and disease severity index, normality of distributions was assessed through qq plots and Shapiro-Wilk tests, and homogeneity of variance was assessed through Levene’s tests. Data for which the null hypothesis of normality was not rejected was analysed through one-way ANOVA data; cases where normality was rejected were analysed with Kruskal-Wallis tests. Distributions with homogenous variances were analysed through Least Significant Differences post-hoc tests. Distributions with non-homogeneous variances were analysed with Dunnett’s post-hoc tests. When data did not support rejection of normality, two-tailed t-tests were used to compare differences between water control and grouped treatments of different ABA concentrations. Experiments were repeated twice with similar results.

## Results

### Data exploration and sorting

A total of 224 data points were collected from primary literature directly reporting resistance phenotypes (Figure 1, Table S1). 194 data points were from studies using Arabidopsis or tomato as the experimental species, representing 58% and 29% of the data, respectively (Figure 1A). The majority of data represented experiments conducted in leaves, with only 14/225 data points in fruit tissue (Figure 1B). There were 55 total different genotypes in this dataset that were categorised by type of mutation into three classes: wildtype (WT), ABA mutant (including signalling and synthesis variants), and ‘Other’ (primarily callose or other hormone-disrupted mutants). ABA mutants represented 46% of the data points (Figure 1C). Regarding plant age, 94% of datapoints were obtained in plants of the age of or younger than 6 weeks (Figure 1D). 23%, 49% and 28% of the data points correspond to biotrophic, necrotrophic and hemibiotrophic pathogens, respectively (Figure 1E). Fungi represent the majority of the data with 66% of the points, while bacteria made up 30% and oomycetes made up 4% (Figure 1F). Concentrations of ABA range from 0 to 100,000uM, although 95% of the data points were between 10 and 750uM (Figure 1G). In this dataset, the level of ‘percentage resistance change’ varied from −800 to 91. 132 data points were labelled ‘Susceptible’ (negative change in resistance) and 93 datapoints ‘Resistant’ (positive change in resistance) (Figure 1H).

### CART models

The CART algorithm was used to produce a decision tree (DT) to unpick which variables are involved in, and the most influential in, determining resistance phenotypes linked to ABA. A DT predicting a binary response of resistance had a predictive accuracy on unseen data of 76.3%. Rpart determined the most important variables in DT (Table 1). ABA concentration (in _μ_M) was determined as the most important variable. Plant age (in weeks), plant species and mutant type were also determined as important predictive variables.

**Table 1:** Importance of variables in DT and RF determined by rpart and RF. Data described in Table S1 and Table S3.

| Model | Variable | Predictive importance |
| --- | --- | --- |
| DT | ABA concentration ($\mu\text{M}$) | 21.7 |
|  | Plant age (weeks) | 13.54 |
|  | Plant mutant | 12.98 |
|  | Plant species | 11.87 |
|  | Pathogen lifestyle | 9.89 |
| RF | ABA concentration ($\mu\text{M}$) | 35.02 |
|  | Plant species | 15.34 |
|  | Pathogen lifestyle | 14.55 |
|  | Plant age (weeks) | 12.7 |

### Random Forest models

A random forest model was created for predicting a binary resistance output. Increasing the number of trees used in the forest increased the predictive accuracy of the model, however beyond 400 trees the predictive capabilities of our model did not improve. The final RF model was therefore generated by using 400 trees. This model had an out-of-bag (OOB) estimate of error rate of 20.98%. The error rate in unseen data for predicting ‘resistance’ was 27%, the error rate for predicting ‘susceptibility’ was 15%. As in the DT model, ABA concentration (_μ_M) was considered the most important variable in the RF model (Table 1). Similar to DT models, RF predicts plant age (in weeks), plant species and pathogen lifestyle as important variables.

The DT (Figure 2) predicted that ABA concentrations above 38_μ_M more frequently lead to susceptibility. This tree predicted that concentrations below 38_μ_M infected with biotrophic pathogens display resistance and that with ABA concentrations below 38_μ_M infected with non-biotrophic pathogens tomato plants display resistance. The DT predicts that ABA concentrations above 38_μ_M infected with non-fungal pathogens display susceptibility. This DT shows the impact of age in resistance: it predicts that with ABA concentrations above 38_μ_M, during fungal infections ‘other’ mutant plants younger than 5.3 weeks display greater resistance than plants over 5.3 weeks.

**Figure 2:**
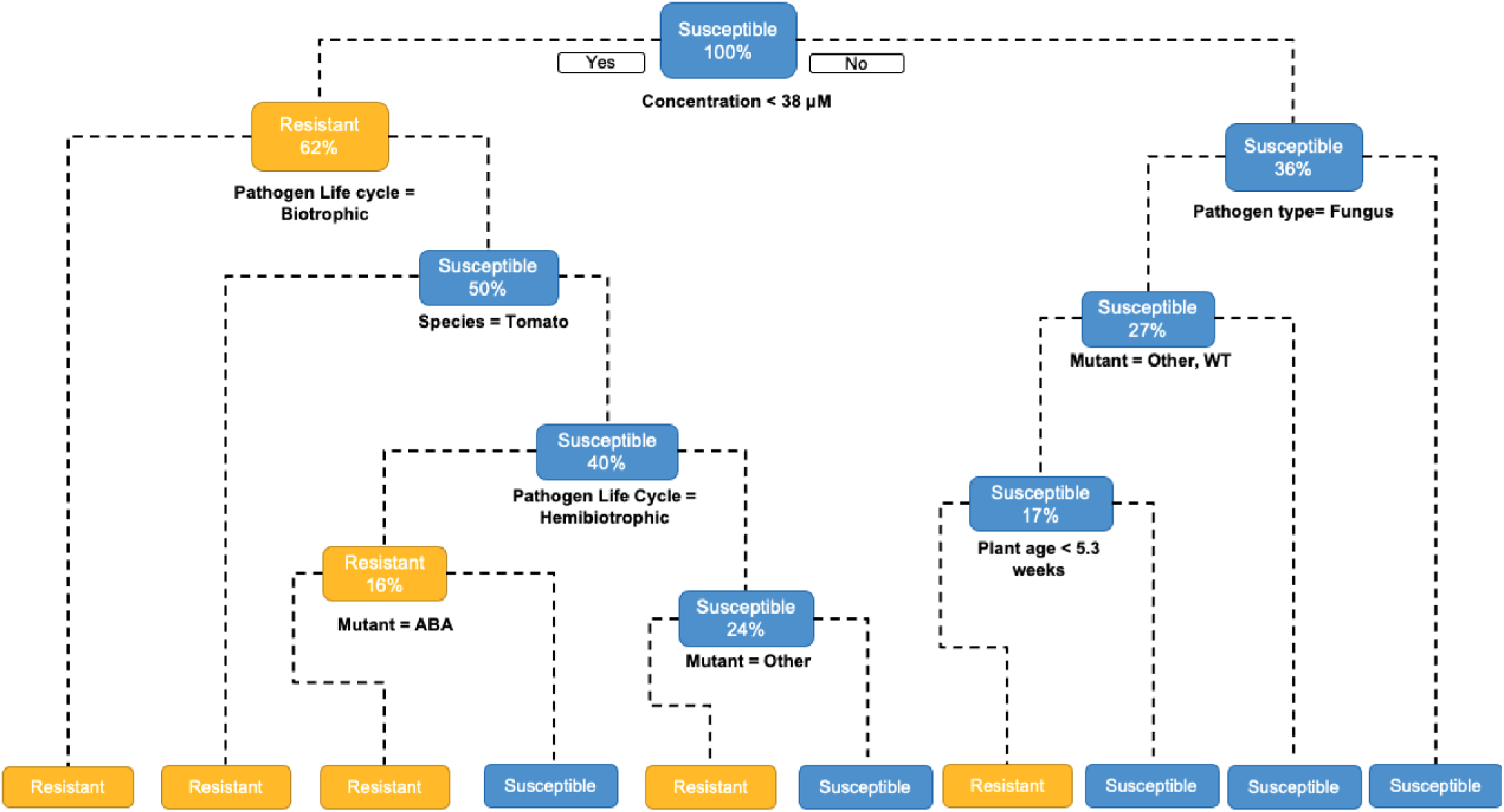
Decision tree (DT) predicting a binary resistance response of ‘Susceptible’ or ‘Resistant’. Yellow boxes indicate resistance and blue boxes indicate susceptibility. Data described in Table S1.

### Effect of ABA on disease resistance against *Botrytis cinerea* and *Phytophthora infestans*

Our RF and DT results highlighted that ABA concentration, pathogen lifestyle and plant age are important variables in resistance predictions. Seeking to refine our quantitative understanding of these influences, we therefore experimentally tested these specific variables. Tomato was used as the experimental plant species as it appears as a split in our decision tree (Figure 2) and is one of the most represented species in our data (Table S1).

The effect of exogenous ABA application at different concentrations on susceptibility against *B. cinerea* was assessed in tomato plants of different ages (i.e. four week and eight-week-old). ABA treatment had a statistically significant effect on the phenotype in leaves from 4-week-old plants against *B. cinerea* (Figure 3A; ANOVA, *p*= 0.04), with ABA generally causing an increase in susceptibility. 50_μ_M.In contrast, in older plants, ABA concentrations of 50_μ_M and above enhanced resistance against *B. cinerea* (Figure 3C; Kruskal-Wallis, *p* = 0.001). The concentration of 100_μ_M led to a significant, dramatic reduction in disease severity (Figure 3C). These results suggest that the effect of ABA on susceptibility against *B. cinerea* is influenced by ABA concentration and the plant age.

**Figure 3:**
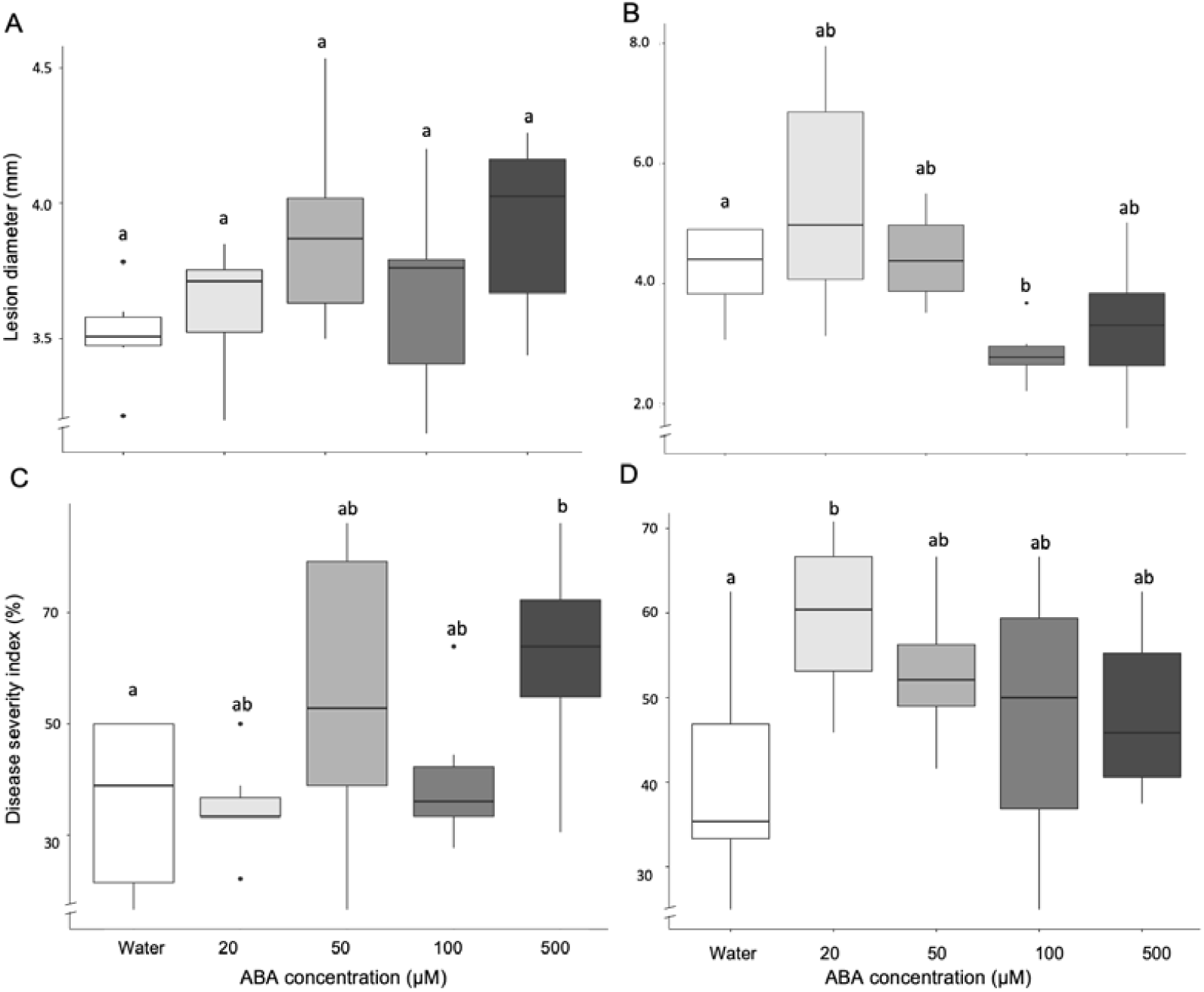

Disease resistance phenoty pes in tomato. Four and eight-week-old tomato leaves were sprayed with 20, 50, 100, 500 _μ_M of ABA 24 hours pre inoculation. (A) Lesion diameters in four-week-old tomato leaves caused by *Botrytis cinerea* at 3 dpi. (B) Lesion diameters in eight-week-old tomato leaves caused by *B. cinerea* at 3 dpi. (C) Disease severity index in four-week-old tomato leaves converted from the percentage of lesions in 6 disease categories at 7dpi with *Phytophthora infestans*. (D) Disease severity index in eight-week-old tomato leaves converted from the percentage of lesions in 6 disease categories at 7dpi with *P. infestans*. Boxes denote 25^th^ and 75^th^ percentile, bars display min and max values. Different letters denote significant differences among treatment groups (LSD for A and D and Dunnets’ for B and C post hoc tests; p < 0,05; n = 6).

The effect of ABA application was also studied against the biotrophic pathogen *P. infestans*. In young leaves, ABA had a statistically significant effect on susceptibility against this pathogen (Figure 3E; ANOVA, *p*= 0.012), with the concentration 500_μ_M causing increased susceptibility in four-week-old leaves (p= 0.029). In older leaves, ABA treatment increased susceptibility in all instances (Figure 3G; ANOVA, *p*= 0.04). However, lower range concentrations had a more severe effect on disease incidence, with 20_μ_M leading to statistically significant increase in susceptibility (*p* = 0.003).

### Decision trees with extended experimental data

Comparing our DT predictions (Figure 2) to our experimental results (Figure 3) highlighted similarities and differences in our model. Our DT predicted only 1/8 of our *B. cinerea* infection results correctly. The prediction for *P. infestans* infected plants was more accurate with 6/8 experimental results being predicted by DT correctly. For a complex question like the role of ABA, such imperfect performance is not surprising. We do not claim to be solving the question of how ABA influences defence in all circumstances, but rather demonstrating a quantitative approach for knowledge curation and generation. To this end, we took the next step in what we propose as an iterative process: refining an initially imperfect model by including observations from new and rationally chosen experiments (Table S2, Table S4). We thus added our experimental findings to our dataset, forming DT2 (Figure 4). The predictive abilities of DT2 remained relatively unchanged, with a predictive accuracy on unseen data of 70%. However, DT2 is now capable of better capturing the new experiments above. For *B. cinerea* infections 6/8 of our experimental results were predicted correctly from DT2, 6/8 were predicted correctly from *P. infestans* infection. The most important variable in DT2 was maintained as ABA concentration. The addition of our experimental data to our model altered some later splits in the DT2 compared to DT1; the split of age was lowered from 5.3 weeks to 4.5 weeks and an additional split was created for >= concentration 375uM, with higher concentrations predicting resistance in all instances.

**Figure 4:**
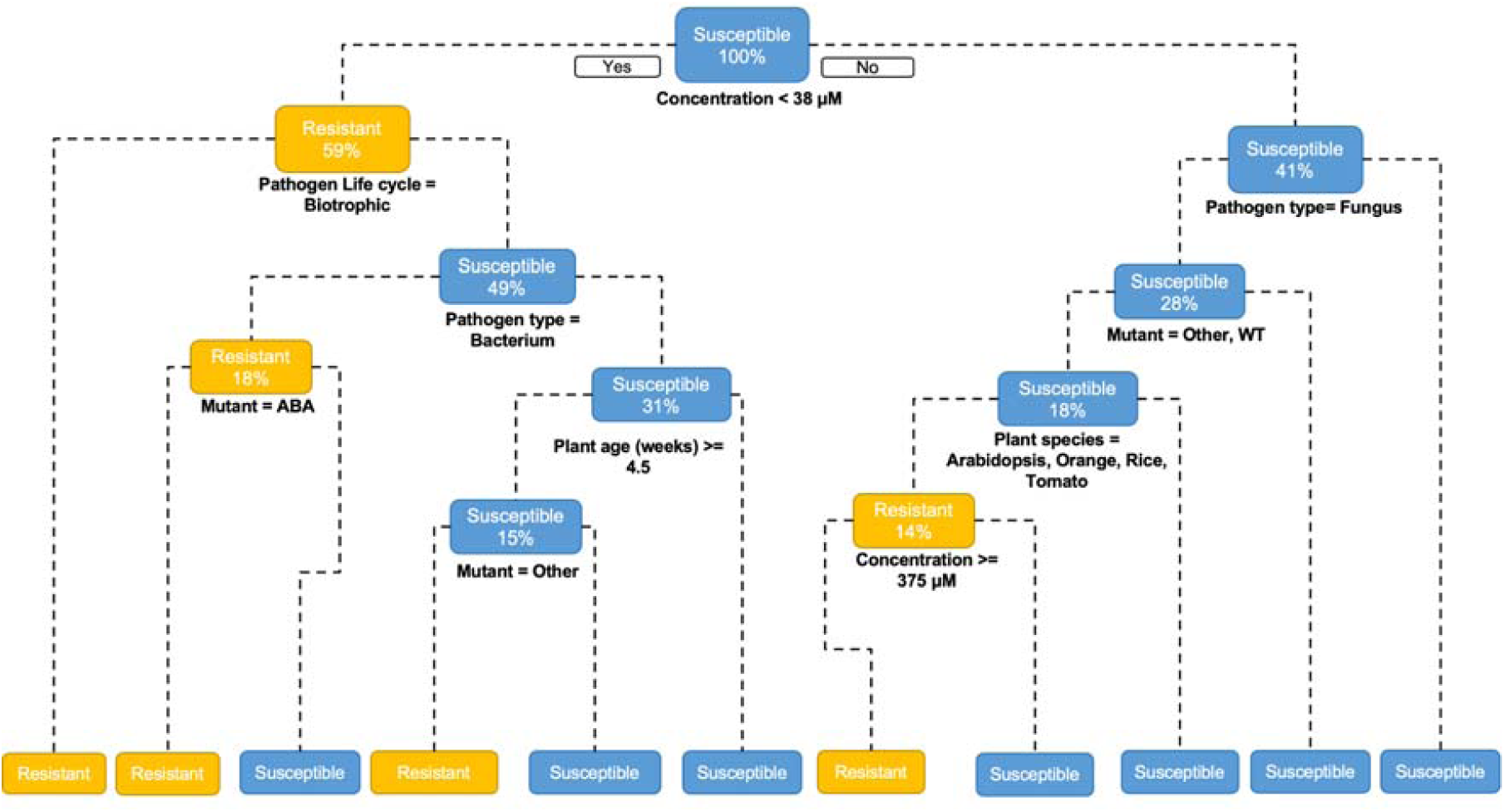

Decision tree (DT) including our experimental data predicting a binary resistance response of ‘Susceptible’ or ‘Resistant’. Yellow boxes indicate resistance and blue boxes indicate susceptibility. Data described in Table S2.

## Discussion

The role of ABA in pathogen defence has remained highly ambiguous (Ton et al., 2009). In this study we used supervised ML techniques to attempt to resolve these relationships and determine the most influential factors in ABA resistance phenotypes. In our predictions and experiments, overall, we can observe pronounced roles for ABA on susceptibility to pathogens. ABA concentration, pathogen lifestyle and plant age were shown to be important influencers of resistance, therefore illustrating ABA’s regulatory role in the activation of major defence pathways and its specificity in individual pathosystems.

### Importance of the concentration

DTs and RF analyses predict the concentration as the primary determinant factor on the role of ABA. Our DT predicts that concentrations higher than 38_μ_M, result in susceptibility (Figure 2). Our experimental work showed in general terms that ABA application at high concentrations is associated with susceptibility (Figure 3). However, our experimental work did not show a consistent pattern of susceptibility as concentration of ABA was increased or as it passed a particular concentration threshold. For instance, we were unable to observe changes at the threshold concentration predicted and instead variable levels of susceptibility were observed (Figure 3). Moreover, whereas increasing concentrations of ABA led to enhanced susceptibility, the concentration of 100_μ_M was extremely effective in triggering resistance against *B. cinerea* (Figure 3C). The differences observed between the predictions and the experimental work may be in part due to differences in the basal levels of ABA in plants. Achuo *et al*., demonstrated major discrepancies in the endogenous ABA levels found in Moneymaker tomato plants. Basal quantities varied within identical experimental set ups from 408 picomol g−1 FW to approximately 2,500 picomol g−1 FW (Achuo et al., 2006). Therefore, basal concentrations of ABA can be different depending on the experimental conditions and could explain discrepancies between predictions and observations.

### Importance of the plant age and pathogen lifestyle

DT analysis allowed us to explore the effect of plant age and pathogen lifestyle. Generally, DT predicted that ABA triggers resistance against biotrophic pathogens with low ABA concentrations. When it comes to the effect against fungal pathogens, DT predicts that ABA affects the resistance phenotype in an age-dependent manner. Our DT predicts susceptibility in older plants, however experimental work has confirmed that ABA triggers resistance to the necrotrophic fungal pathogen *B. cinerea* only in older leaves.

The reasons behind the enhanced resistance observed after ABA treatments in older plants may be due to age related resistance (ARR). It is well reported that plant age influences pathogen resistance, with young juvenile leaves displaying enhanced susceptibility (Kus et al., 2002). This is often due to important defence functions, such as PR gene expression, being upregulated in older leaves (Kus et al., 2002, Li et al., 2020). Moreover, different developmental stages, such as the transition to floral stage, are associated with increased resistance in a range of species including Arabidopsis, tobacco and tomato (Develey-Rivière and Galiana, 2007, Hu and Yang, 2019). This resistance has been linked to a positive correlation between development and defence-related processes such as phytoalexin production and resistance gene expression (Bell, 1969, Li et al., 2020). The DT model produced in this study however did not predict this ARR. This is likely due to the small age interval of our data, as 3-6 weeks seems to be the most popular age range for infection studies (Figure 1, Table S1). Furthermore, a limitation of our data collection could also be responsible for the lack of ARR in the DT predictions, mostly based on a threshold of age. Whereas we have recorded and assumed that below ∼four weeks old is considered “young” and ∼eight weeks is considered “old”, this may not be true for all plant species studied. For instance, a four-week-old Arabidopsis plant may no longer be considered young, and an eight-week-old orange tree cannot be considered old. A more accurate assessment of ARR would therefore come from the analysis of the impact of ABA in resistance at different developmental stages for each plant species.

Our results show that pathogen lifestyle determines the effect of ABA in resistance; this can be linked to the modulation of major defence networks associated with the infection strategies. It is well characterised that whereas biotrophic pathogens result in the activation of SA-dependent defences (Mohr and Cahill, 2007), the plant activates JA and ET-dependent defences in response to necrotrophic pathogens (McDowell and Dangl, 2000). Moreover, there is a comprehensively understood antagonistic interaction between SA and JA (Anderson et al., 2004, De Torres Zabala et al., 2009). It is known that *B. cinerea* produces a virulence factor, EPS, that directly modulates the JA/SA antagonism. EPS suppresses JA defences, reduces *proteinase inhibitor I* and *II* (*PI I* and *PI II*) expression and leads to accumulation of SA and NPR1 expression (El Oirdi et al., 2011). Tomato hosts resistance to *P. infestans* through production of defence proteins that have been shown to be ET and SA dependent (Smart et al., 2003). Importantly, it is also known that there are multiple antagonistic responses between ABA and SA, JA, and ET. Therefore, it is easy to speculate that the effect of ABA in resistance against pathogens with different lifestyles is the result of ABA impacting the activation of those hormone-dependent signalling pathways. To our knowledge this is the first example of the impact of exogenous ABA application on tomato resistance against the biotroph *P. infestans*. Previous studies have been carried out on potato slices against this pathogen demonstrating that ABA treatments increase susceptibility (Henfling et al., 1980, Liu et al., 2020). Our experimental work also demonstrates that treatments with 500_μ_M ABA result in susceptibility against *P. infestans* (Figure 3E). Therefore, it is likely that this phenotype is due to disruption of SA defence signalling by ABA. Future studies analysing both genotypes altered in SA and JA signalling will help confirm if ABA application is contributing to susceptibility due to the direct activation or repression of these essential defence pathways.

### Application of machine learning tools to complex biological mechanisms

We suggest that tools from ML (here particularly decision tree and random forest classification approaches) provide an approach for further clarification of the role of ABA. This approach is, unsurprisingly, limited by the amount of data available -- as demonstrated by our original decision tree, which had reasonable predictive performance on test data but failed to capture behaviour in several new experiments. For instance, DT predicts resistance in 4-week-old leaves against *B. cinerea* (Figure 2), however susceptibility is observed experimentally (Figure 3). However, these approaches are not proposed as an immediate solution, but rather as a framework for iteratively refining and improving our understanding. Hence, in an attempt to improve the prediction of the initial data, we added our experimental data into the model (Figure 4). This improved our DT predictions in the tomato-*B. cinerea* pathosystem, increasing correct predictions from 1/8 to 6/8. Therefore, the addition of only 16 new data points (i.e., 7.1% change in the original data) led to changes in DT predictions and far greater accuracy against our own experimental results. This improvement points to the importance of adding under-represented data to the model. Here, the capacity for improvement can likely be attributed to the limited current dataset, in particular the narrow age range represented in tomato with no data points included in leaves from plants older than 5.5 weeks. We believe that these ML approaches can both help unify and interpret observations from heterogeneous studies and to identify relevant characteristics in the role of ABA in defence. As more data is added, particularly on sparsely represented cases and surrounding key tree splitting points, further clarity and prediction refinement will be achieved. Thus, we invite researchers to input their results into the model, especially from data in under-represented groups, to improve its prediction capacity. This could, for example, determine whether our findings of the age-dependent effect of ABA against necrotrophs is specific to tomato or can be extrapolated to other plant species.

### Moving forward with experimental work on the role of ABA in defence

Due to its essential role in abiotic defences, ABA signalling is a common target for engineering stress tolerant plants. Therefore, greater understanding of its controversial role in biotic defences is essential to ensure phenotypes are resistant to both abiotic stress and pathogen challenge. Methodologies used in ABA research vary significantly, this is testament to the many functions of ABA and physiological processes in which it is involved. For example, data examined from literature in this study included repeated ABA spray application, root soaking, biosynthesis mutants, signalling mutants and salt stress treatment, all of which will lead to variable contributions to resistance (Achuo et al., 2006, Audenaert et al., 2002, Song et al., 2011). Despite these, we have been able to identify key characteristics that could determine phenotypes. Whereas it is clear that ABA triggers susceptibility against biotrophic pathogens in a concentration dependent manner, this is not as straightforward against necrotrophic pathogens, where plant age and specific concentrations are relevant. Importantly, it is important to understand the function of hormone crosstalk in evolution and survival, as they are an essential characteristic to prevent singular resource allocation against a single stressor (Vos et al., 2015). ABA has been demonstrated to be fully involved in the expression of priming of defence, which does not result in the direct activation of defence mechanisms but the fine-tuning of the response upon subsequent challenges. Due to the key role of ABA in defence crosstalk and priming, it could be hypothesised that ABA plays a central role in buffering any single effort, contributing to noisiness, and therefore playing a central role in the evolution of defence strategies. Therefore, this study can serve as a tool to test that hypothesis and to guide future studies involving the role of ABA in disease resistance.

## Data and code availability

All data and code is publicly available at https://github.com/PlantPriming/ABAresistance

## Acknowledgements

The authors thank Ana Pineda for her support with the first draft of this paper through the scientific writing online course “I focus and write”.

